# RNA sequencing of the i*n vivo* human herpesvirus 6B transcriptome to identify targets for clinical assays distinguishing between latent and active infections

**DOI:** 10.1101/397679

**Authors:** Joshua A. Hill, Minako Ikoma, Danielle M. Zerr, Ryan S. Basom, Vikas Peddu, Meei-Li Huang, Ruth Hall Sedlak, Keith R. Jerome, Michael Boeckh, Serge Barcy

## Abstract

Human herpesvirus 6B (HHV-6B) DNA is frequently detected in human samples, especially after hematopoietic cell transplantation (HCT). Diagnostic assays distinguishing HHV-6B reactivation from latency are limited, and this has contributed to confusion in research and made the design of clinical approaches to diagnose and treat HHV-6-associated diseases challenging. We used RNA sequencing to characterize and compare the HHV-6B transcriptome in multiple *in vivo* and *in vitro* sample types, including 1) whole blood from HCT recipients with and without HHV-6B plasma viremia; 2) tumor tissue samples from subjects with large B cell lymphoma infected with HHV-6B; 3) lymphoblastoid cell lines from subjects with inherited chromosomally integrated HHV-6B or latent infection with HHV-6B; and 4) HHV-6B Z29 infected SupT1 CD4+ T cells. We demonstrated substantial overlap in the HHV-6B transcriptome observed in *in vivo* and *in vitro* samples, although there was variability in the breadth and quantity of gene expression across samples. No HHV-6B transcripts were detected in whole blood samples from subjects without plasma HHV-6B viremia. The HHV-6B viral polymerase gene U38 was the only HHV-6B transcript detected in all RNA-seq data sets and was one of the most highly expressed genes. Using a novel reverse transcription PCR assay targeting HHV-6B U38, we identified U38 messenger RNA in all tested whole blood samples from patients with concurrent HHV-6B viremia, indicating its utility as a diagnostic assay for HHV-6B replication. This study demonstrates the feasibility of pathogen transcriptome analyses in HCT recipients to identify better targets for diagnostic, and potentially therapeutic, applications.

**IMPORTANCE:** Infection with human herpesvirus 6B (HHV-6B), a DNA virus, occurs early in life, results in chronic viral latency in diverse cell types, and affects the population at large. Additionally, HHV- 6B can integrate into germline chromosomes, resulting in individuals with viral DNA in every nucleated cell. Given that PCR to detect viral DNA is the mainstay for diagnosing HHV-6B infection, the characteristics of HHV-6B infection complicate efforts to distinguish between latent and active viral infection, particularly in immunocompromised patients who have frequent HHV- 6B reactivation. In this study, we used RNA sequencing to characterize the HHV-6B gene expression profile in multiple sample types, and our findings identified evidence-based targets for diagnostic tests that distinguish between latent and active viral infection.

## INTRODUCTION

Human herpesvirus 6B (HHV-6B) is a double-stranded DNA virus that infects the majority of people during infancy and establishes latency in a diverse array of cell types.^1^ Immunosuppression due to allogeneic hematopoietic cell transplantation (HCT) results in frequent viral reactivation with detection of HHV-6B DNA viremia in approximately 40-60% of patients. HHV-6B reactivation is the most frequent cause of encephalitis in HCT recipients, affecting 1-10% of patients,^2^ and HHV-6B is associated with a variety of disease processes in both immunocompromised and immunocompetent individuals.^1^ Quantitative polymerase chain reaction (qPCR) assays to detect HHV-6B genomic DNA in biologic samples are limited by the detection of latent virus and low specificity for end-organ disease.^3–7^ This is especially problematic in patients with inherited chromosomally integrated HHV-6B (iciHHV-6B), who have a high burden of cell-associated HHV-6 DNA due to integration of the viral genome in every nucleated cell.^8,9^ These limitations in identification of HHV-6B reactivation and associated pathogenicity confound research and treatment efforts for HHV-6B.^10^

Molecular detection of HHV-6B gene transcripts using reverse transcription real-time qPCR (RT-qPCR) has potential to improve the specificity of diagnostic studies for HHV-6B reactivation. This method of amplifying messenger (m)RNA from infected cells may provide a better approach to distinguish active from latent infections. A few studies have reported the development and application of assays to detect HHV-6 mRNA transcripts including open reading frames (ORFs) U12, U16/17, U60/66, U79/U80, U89/90, and U100.^11–17^ However, these studies have been limited by reliance on *in vitro* data for transcript selection, use of selected mRNA targets based on spliced gene products, and suboptimal RNA preservation due to sample collection methods. High throughput qualitative profiling of transcript changes using microarrays have been used to identify HHV-6B gene expression with *in vitro* cell lines in one study,^18^ but these methods may also produce biased results through the use of predefined probe sets. To our knowledge, only one study performed HHV-6B transcriptome analysis with unbiased next generation RNA sequencing (RNA-seq) in tissue samples from two patients with diffuse large B-cell lymphoma (DLBCL).^19^

We sought to characterize *in vivo* HHV-6B gene expression using high-throughput RNA-seq on polyA-selected RNA isolated from whole blood samples in HCT recipients with HHV-6B reactivation. To facilitate identification of highly expressed mRNA transcripts in biologic specimens to guide development of diagnostic assays, we compared HHV-6B gene expression patterns in whole blood to data from a study using tumor tissue samples and to cell cultures with lytic, latent, and iciHHV-6B. We also assessed whether we could identify distinct host gene transcription and cytokine profiles among HCT recipients with and without HHV-6B DNA detection in plasma samples.

## METHODS AND MATERIALS

### Patients

We identified a cohort of patients who had whole blood collected in PAXgene Blood RNA tubes (PAXgene tubes; QIAGEN, Germantown, MD) within the first six weeks of autologous or allogeneic HCT as part of an unrelated study. We then identified plasma samples from these subjects that were collected on the day of PAXgene tube collection and temporally proximal, both before and after. We tested these samples for HHV-6B DNA using qPCR. Patients who had plasma HHV-6B DNA detected within 7 days of PAXgene tube collection were selected for this study and were categorized as cases; one patient was included with 11 days between sample collection and HHV-6B detection (**Table 1**). Patients without HHV-6B detection in plasma samples within 7 days of PAXgene tube acquisition were used as controls and matched in a 1 to 1 ratio to cases based on the day post-HCT of PAXgene tube collection, age, race, sex, type of HCT (autologous versus allogeneic), donor relation, human leukocyte antigen (HLA) matching, neutrophil engraftment status, and acute graft-versus-host disease (aGVHD) status (**Table 1**).

**Table 1.** Post-hematopoietic cell transplant day of HHV-6B plasma detection relative to day of whole blood PAXgene tube collection.

| **Subjects** | **PAXgene day<sup>a</sup>** | **Closest plasma HHV-6B detection day<sup>a</sup>** | **Closest HHV-6B plasma viral load<sup>b</sup>** |
| --- | --- | --- | --- |
| Case 1 | 26 | 20 | 6,900 |
| Case 2 | 22 | 22 | 2,000 |
| Case 3 | 17 | 21 | 1,567 |
| Case 4 | 31 | 27 | 518 |
| Case 5 | 7 | 18 | 2,012 |
| Case 6 | 13 | 17 | 4,827 |
| Case 7 | 26 | 25<sup>c</sup> | 4,057 |

<sup>a</sup>Day represents the number of days post-HCT.
<sup>b</sup>Data are presented in units of copies/mL.
<sup>c</sup>This subject also had HHV-6B DNA detected on day 32 (+6) with a viral load of 364 copies/mL.

We also used data from two patients with DLBCL from a published study in which RNA-seq on tumor tissue samples demonstrated a broad profile of HHV-6B transcripts consistent with lytic transcription.^19^

### Samples

PAXgene tubes were filled with 2.5 ml of whole blood obtained directly from HCT recipients, and samples were stored according to the manufacturer’s instructions. Plasma samples were leftover clinical specimens used for cytomegalovirus testing. For *in vitro* comparisons, we obtained a cell culture of HHV-6B Z29 infected SupT1 CD4+ T cells (provided by the HHV-6 Foundation). Cells were harvested 48 hours after infection. We also obtained two beta lymphoblastoid cell lines (B-LCLs) derived from pre-HCT PBMCs from one subject with low-level virus detection due to acquired, latent HHV-6B infection and one subject with iciHHV- 6B identified in a prior study.^20^ The presence or absence of iciHHV-6B in each of the B-LCLs was confirmed by droplet digital PCR as previously described.^9^

### HHV-6B DNA testing

We tested plasma samples for HHV-6B DNA using high-throughput real-time qPCR that detects 1 copy of HHV-6 DNA per reaction (25 copies/ml) and distinguishes between species A and B as previously described.^21^

### RNA extraction

We isolated and extracted total RNA from PAXgene tubes according to the manufacturer’s instructions using the MagMAX Blood RNA kit (Ambion, Foster City, CA). RNA quality was determined by electropherogram (showing 28S, 18S, and 5S bands) and RNA integrity number using an Agilent 2200 Bioanalyzer (Agilent Technologies, Santa Clara, CA). Total RNA concentration was obtained from an absorbance ratio at 260 and 280 nm using a NanoDrop ND-2000 spectrometry instrument (NanoDrop Technologies, Wilmingtion, DE). Since whole blood RNA consists mostly of globin mRNA, we performed globin depletion using the GLOBINclear kit (Ambion, Foster City, CA) to maximize detection of less abundant mRNA and further purified and concentrated RNA on appropriate columns before RNA library preparation.

### RNA sequencing

We performed polyA-RNA (coding mRNA) selection to minimize DNA contamination. RNA-seq libraries were prepared using the Ovation Single Cell RNA-seq System (NuGen Technologies, Inc., San Carlos, CA, USA). Library size distributions were validated using an Agilent 2200 TapeStation (Agilent Technologies, Santa Clara, CA, USA). Additional library quality control, blending of pooled indexed libraries, and cluster optimization were performed using Life Technologies’ Invitrogen Qubit^®^ 2.0 Fluorometer (Life Technologies-Invitrogen, Carlsbad, CA, USA). Barcoded RNA-seq libraries were run individually on a flow cell lane using an Illumina cBot for whole blood from samples HCT recipients with HHV-6B viremia (cases) and the SupT1 CD4+ cell culture; whole blood from HCT recipients without HHV-6B viremia (controls) and LCLs were duplexed or triplexed. Sequencing was performed using an Illumina HiSeq 2500 in rapid-mode employing a paired-end, 50-base read length sequencing strategy. Image analysis and base calling were performed using Illumina’s Real Time Analysis v1.18 software, followed by ‘demultiplexing’ of indexed reads and generation of FASTQ files using Illumina’s bcl2fastq Conversion Software (v1.8.4). Reads that passed the quality check were aligned to a HHV-6B reference genome (Genbank accession number NC_000898) using TopHat (v2.09) along with Bowtie 2 (v2.2.3).^22,23^ Aligned reads obtained from TopHat were visualized using Geneious software (Biomatters Limited) to assess the extent and depth of coverage, sequence alignments, and identified splice junctions. Read counts per HHV-6B gene enumeration were generated using HTseq-count (v0.6.1), employing the “intersection-strict” overlap mode.^24^

### ORF U38 RT-qPCR assay development

RNA was extracted from 2.5 ml of whole blood collected in PAXgene tubes using the PAXgene blood RNA kit (Qiagen, Germantown, MD) and eluted into 80 µl of BR5 buffer. To prevent amplification of HHV-6B genomic DNA, samples were treated with DNAse as part of the extraction protocol. We used primers (forward: TGCCCGATTYTGAAAAAGCT, reverse: CCTGTGGGTATTCATAAAATTTTG) and probe (FAM-CTCCCGCGCTTTGCACAGACG-BHQ) specific to a conserved region of ORF U38 using Primer Express 3.0.1 (Applied Biosystems, Foster City, CA). One step RT-PCR was performed using the QuantiTect Probe RT-PCR kit (Qiagen, Germantown, MD) and the 7500 fast real time PCR system. Each 25 µl RT-qPCR reaction contained 0.25 µl QuantiTect virus RT mix (100x), 5 µl QuantiTect virus master mix (5x), 0.5 µl ROX, 0.4 µM primers and 0.2 µM probe, 10 µl RNA, and water to volume. The thermocycling conditions were as follows: 20 minutes at 50° C for reverse transcription, 5 minutes at 95° C for PCR activation, 45 cycles at 95° C for 15 seconds followed by 45 seconds at 60° C. The assay was first verified using standards created from primer-specific ORF U38 transcripts isolated from an HHV-6B infected cell line. An RT-qPCR reaction excluding reverse transcriptase was performed on all standards and samples to assess for DNA contamination.

### Statistics

To assess global differences in HHV-6B gene expression profiles from different sample types, we performed hierarchical clustering analysis using Clustered Image Maps software (CIMminer). The Manhattan distance matrix was computed based on the respective abundance of each viral gene detected in each patient. To allow comparisons between datasets obtained from different sample types, read counts per HHV-6B gene were normalized by calculating viral reads per million unique mapped genomic reads (VPMM). These values were used as input for hierarchical clustering using the complete linkage-clustering algorithm. Data sets representing viral gene expression profiles were visualized by heat maps.

We performed a differential gene expression analysis of host cellular gene expression profiles comparing normalized read counts from RNA-seq data sets between cases and controls. Paired analysis was performed using the Bioconductor package DESeq22 software.^25^

### Sequencing data accession number

RNA-seq generated from our samples are available at the Gene Expression Omnibus (GEO) website under accession number: GSE115533. RNA-seq data sets for two patients with DLBCL (SRS405408 and SRS405443) were downloaded from the NIH database of genotypes and phenotypes (dbGap; http://www.ncbi.nlm.nih.gov/sites/entrez?db=gap) using accession code phs000235.v2.p1. These RNA-seq data sets are freely available through controlled access.

## RESULTS

### Patients

We identified seven subjects (cases 1-7) with whole blood PAXgene tubes obtained post-HCT who had concurrent HHV-6B DNA detection in plasma by qPCR with a median of 4 days (range, 0-11) between plasma detection and PAXgene tube collection. HHV-6B was detected in plasma samples prior to, on the day of, and after the day PAXgene tube collection in 2, 1, and 3 cases, respectively; HHV-6B was detected prior to and after the day of PAXgene tube collection in 1 case (**Table 1**). Matched controls who had a PAXgene tube collected and did not have HHV-6B DNA plasma detection within 7 days of the PAXgene tube collection were identified for cases 1-5. Demographics and clinical characteristics of the cohort are presented in **Table 2**. None of the subjects developed HHV-6-associated end-organ disease. In addition, we obtained RNA-seq data sets from two patients with DLBCL who had HHV-6B gene expression demonstrated in tumor tissue samples.^19^

**Table 2.** Demographic and clinical characteristics of post-hematopoietic cell transplant cases (with HHV-6B plasma detection) and controls (without HHV-6B plasma detection) with whole blood collected in PAXgene tubes. HLA indicates human leukocyte antigen; GVHD, graft-versus-host disease; AML, acute myelogenous leukuemia; NHL, non-Hodgkin’s lymphoma; MDS, myelodysplastic syndrome; MM, multiple myeloma; HD, Hodgkin’s disease.

| Patients | Age | Sex | Race | Disease | Donor | HLA-match<sup>a</sup> | Engraftment day<sup>c,d</sup> | GVHD day<sup>c,e</sup> |
| --- | --- | --- | --- | --- | --- | --- | --- | --- |
| Case 1 | 46 | Male | Caucasian | AML | Unrelated | Mismatched<sup>b</sup> | 17 | - |
| Control 1 | 56 | Male | Hispanic | AML | Unrelated | Mismatched<sup>b</sup> | 8 | 47 |
| Case 2 | 59 | Male | Caucasian | NHL | Related | Matched | 15 | 27 |
| Control 2 | 64 | Male | Caucasian | MDS | Unrelated | Matched | 27 | 49 |
| Case 3 | 54 | Female | Caucasian | MM | Unrelated | Matched | 21 | 30 |
| Control 3 | 57 | Female | Caucasian | NHL | Unrelated | Matched | 21 | - |
| Case 4 | 53 | Female | Caucasian | MM | Related | Matched | 13 | - |
| Control 4 | 57 | Female | Caucasian | MDS | Related | Matched | 20 | - |
| Case 5 | 61 | Male | Caucasian | MM | Self | Matched | 16 | - |
| Control 5 | 72 | Male | Caucasian | MM | Self | Matched | 16 | - |
| Case 6 | 44 | Male | Caucasian | HD | Unrelated | Matched | 12 | 28 |
| Case 7 | 54 | Female | Caucasian | AML | Unrelated | Mismatched<sup>b</sup> | 20 | 39 |

<sup>a</sup>Matched indicates 10/10 allele or antigen matching.
<sup>b</sup>Umbilical cord blood was used as the source of donor cells.
<sup>c</sup>Day represents the number of days post-HCT.
<sup>d</sup>Engraftment was defined as the first of three consecutive days of an absolute neutrophil count >500 cells/ $\mu$ L.
<sup>e</sup>GVHD was defined as previously described in Leisenring et al, Blood 2006;108:749-755.

### mRNA isolation from PAXgene tubes

We isolated sufficient mRNA to allow for high throughput sequencing for five of seven cases and three of five controls (**Table 3**). There was substantial variability in complete blood counts for each subject at the time of PAXgene tube collection due to sample acquisition within the first few weeks after HCT (**Table 3**). The lowest yields of total RNA were observed for individuals with the lowest absolute white blood cell (WBC) counts. The two cases and two controls with insufficient RNA recovery had PAXgene tubes obtained the earliest after HCT (within 14 days) and prior to neutrophil engraftment.

**Table 3.** Total WBC count (for whole blood samples), mRNA isolation, and HHV-6B gene transcript counts in post-HCT whole blood samples, DLBCL tissue samples, and SupT1 CD4+ T cells with lytic HHV-6B Z29 infection. WBC indicates white blood cell; mRNA, messenger RNA; VPMM, viral reads per million unique mapped genomic reads; DLBCL, diffuse large B cell lymphoma; NA, not applicable; ND, not done.

| Sample Type | Total WBC<sup>a</sup><br>(cells/mm<sup>3</sup>) | Total mRNA (ng) | Total HHV-6B<br>VPMM |
| --- | --- | --- | --- |
| **Whole blood** |  |  |  |
| Case 1 | 7.0 | 5,075 | 0.02<sup>b</sup> |
| Case 2 | 5.5 | 2,523 | 14 |
| Case 3 | 2.1 | 6,687 | 0.03<sup>b</sup> |
| Case 4 | 3.7 | 9,529 | 0 |
| Case 5 | 0.06 | Insufficient | ND |
| Case 6 | 1.6 | Insufficient | ND |
| Case 7 | 5.2 | 3,973 | 19 |
| Control 1 | 3.0 | 9,222 | 0 |
| Control 2 | 1.3 | 6,467 | 0 |
| Control 3 | 0.3 | Insufficient | ND |
| Control 4 | 9.3 | 12,093 | 0 |
| Control 5 | 0.2 | Insufficient | ND |
| **DLBCL tissue** |  |  |  |
| Case 1 | NA | ND | 21 |
| Case 2 | NA | ND | 94 |
| **SupT1 CD4+ T cells** | NA | ND | 7,040 |

<sup>a</sup>WBC counts were drawn on the same day as the PAXgene tube.
<sup>b</sup>U38 was the only transcript detected.

### Comparison of the HHV-6B transcriptome in whole blood, DLBCL tumor tissue, and SupT1 CD4+ T cells with lytic HHV-6B Z29 infection

We sought to characterize and compare the relative abundance of HHV-6B transcripts between whole blood from post-HCT patients with and without HHV-6B plasma viremia, tissue samples from patients with DLBCL, and a SupT1 CD4+ T cell culture with lytic HHV-6B Z29 infection. In post-HCT whole blood samples from the five cases and three controls with sufficient RNA isolation, transcriptome analysis using RNA-seq identified HHV-6B gene expression in four of five cases and zero of three controls (**Table 3**). Overall, whole blood samples had low HHV- 6B gene counts with a range of 0.03 to 19 VPMM per sample (**Table 3**). In comparison, tumor tissue samples from two patients with DLBCL had 21 and 94 HHV-6B VPMM (**Table 3**). The *in vitro* SupT1 CD4+ T cell culture with lytic HHV-6B Z29 infection had the highest HHV-6B gene count with 7,040 VPPM (**Table 3**).

In post-HCT whole blood, a broad array of HHV-6B genes was detected from cases 2 and 7 including immediate early, early, and late genes (**Tables S1, S2; Figure 1**). In cases 1 and 3, we only detected HHV-6B reads mapping to ORF U38. Tumor tissue samples from the two patients with DLBCL had an overall increased diversity of HHV-6B gene expression to most HHV-6B genes. The *in vitro* SupT1 CD4+ T cell culture with lytic HHV-6B infection demonstrated the greatest diversity of HHV-6 gene expression. Transcripts for U13 and the B genes (B1, B2, B3, B4, B5, B8, B9) were only detected in the HHV-6B infected SupT1 CD4+ T cell culture. The complete data sets of absolute and normalized viral reads are shown in **Tables S1 and S2**.

**Figure 1:**
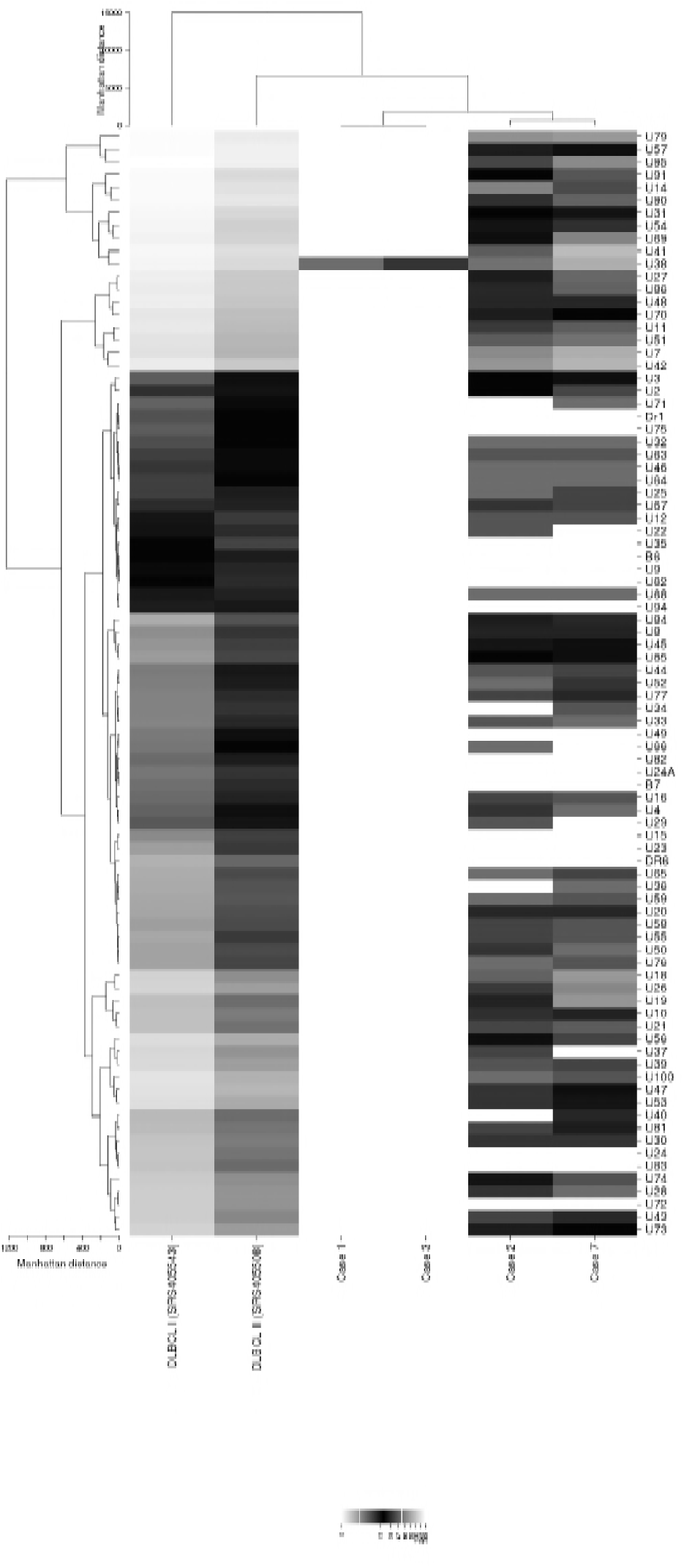
Heat map of hierarchical clustering analysis of the HHV-6B transcriptome from whole blood and DLBCL tumor samples. We performed hierarchical clustering analysis to compare the normalized read numbers mapped to each HHV- 6B gene for whole blood and tumor tissue samples. Hierarchically clustered viral genes (rows) from each sample type (columns) are shown in dendrograms and demonstrate differential clustering. Cases 1, 2, 3, and 7 are from whole blood samples in subjects with HHV-6B DNA detection in plasma after allogeneic HCT. DLBCL I and II are tumor tissue samples from two individuals with DLBCL. DLBCL I was co-infected with Epstein-Barr virus.

We performed hierarchical clustering analysis to compare the normalized read numbers mapped to each HHV-6B gene from whole blood and tumor tissue samples, and the results are displayed in the form of a heat map in **Figure 1**. In this analysis, whole blood samples clustered together with one case of DLBCL. The other case of DLBCL clustered separately indicating a distinct viral gene expression pattern. Results from the SupT1 CD4+ T cell culture could not be included in this figure due to exponentially higher gene expression.

We compared the top ten HHV-6B transcripts with the highest normalized number of reads from each sample type (**Table 4)**. This demonstrated that reads mapping to HHV-6B transcripts associated with expression of ORF U38, a viral polymerase gene, were the only transcripts detected at relatively high levels in all the RNA-seq data sets. Overall, five of the top ten HHV-6B genes (ORFs U95, U79, U41, U38, and U14) were shared between all three sample types, demonstrating high concordance between *in vitro* and *in vivo* HHV-6B gene transcripts in the setting of lytic infection. Reads mapping to late gene transcripts (ORFs U57, U54, U41, and U38) were more prevalent in the DLBCL tissue samples than whole blood samples (ORF U38 only).

**Table 4.** The top ten most highly expressed HHV-6B gene transcripts detected in post-HCT whole blood samples, DLBCL tissue samples, and SupT1 CD4+ T cells with lytic HHV-6B Z29 infection. DLBCL indicates diffuse large B cell lymphoma; VPMM, viral reads per million unique mapped genomic reads; E, early; IE, immediate early; L, late. Bolded genes are those that were in the top ten most highly expressed transcripts for all sample types. We only used one SupT1 CD4+ T cell culture. The temporal pattern of expression is defined as previously described.^33^

| Whole blood (n = 2)<sup>a</sup> |  | DLBCL (n = 2) |  | SupT1 CD4+ T cells (n = 1) |  |
| --- | --- | --- | --- | --- | --- |
| Gene | Average VPMM<sup>b</sup> | Gene | Average VPMM<sup>b</sup> | Gene | VPMM |
| U42 (E) | 0.5 | **U95** (IE) | 2.4 | **U41** (E) | 325.0 |
| U7 (E) | 0.5 | U57 (L) | 2.3 | U11 (L) | 286.0 |
| **U41** (E) | 0.5 | **U79** (IE) | 2.1 | **U14** (E) | 285.9 |
| **U38** (L) | 0.4 | U90 (E) | 1.8 | **U38** (L) | 269.4 |
| **U79** (IE) | 0.4 | **U14** (E) | 1.7 | U100 (L) | 264.2 |
| U18 (E) | 0.4 | U91 (E) | 1.7 | U57 (L) | 243.4 |
| **U95** (IE) | 0.3 | **U41** (E) | 1.5 | U86 (IE) | 222.3 |
| **U14** (E) | 0.3 | U31 (L) | 1.4 | **U95** (IE) | 222.1 |
| U51 (E) | 0.3 | **U38** (L) | 1.4 | U54 (L) | 211.5 |
| U19 (E) | 0.3 | U54 (L) | 1.3 | **U79** (IE) | 207.7 |

<sup>a</sup>The data sets obtained from cases 1 and 3 were omitted due to a low number of reads to only one transcript (U38).
<sup>b</sup>Normalized read counts expressed in VPMM were averaged between samples.

### HHV-6B transcriptome in B-LCLs with latent and iciHHV-6B

We did not detect reads aligning to any specific viral transcripts in data sets from a B-LCL sample with latent HHV-6B infection from post-natal infection or a B-LCL sample with inherited chromosomally integrated HHV-6B. Although we did identify several reads containing the TAACCC nucleotide sequence, we could not determine the origin of this RNA transcript given that this telomeric repeat sequence is found in both HHV-6B and human genomes.

### Development and validation of a RT-qPCR assay for ORF U38

We selected ORF U38 as the best candidate for a diagnostic target to identify lytic HHV-6B replication in biologic samples based on our findings that ORF U38 was highly expressed in all sample types and the only transcript detected in all samples with HHV-6B gene expression. To confirm the detection of reads mapping to ORF U38, we developed a specific RT-qPCR assay to detect this mRNA transcript. Using this newly developed assay, we tested mRNA extracted from whole blood in four of the post-HCT cases with HHV-6B plasma detection who had a second available PAXgene tube. We detected the ORF U38 transcript in all four cases (**Table 5**). Additionally, the high sensitivity of this assay was demonstrated by the detection of U38 mRNA from one subject in which RNA-seq did not identify any HHV-6B gene transcripts (case 4), as well as another case in which there was an insufficient amount of isolated RNA to perform RNA-seq (case 5). The number of ORF U38 copies detected by RT-qPCR correlated with the number of reads mapping to ORF U38 by RNA-seq (*r*^2^ = 1.0; *p* = 0.08), but this did not reach statistical significance, likely due to the small sample size.

**Table 5.** HHV-6B ORF U38 detection by RNA-seq and RT-qPCR in post-HCT whole blood samples. ND indicates not done.

| Subjects | U38 reads,<br>RNA-seq | U38 RNA copies/mL,<br>RT-qPCR | U38 DNA copies/ml,<br>qPCR<sup>a</sup> |
| --- | --- | --- | --- |
| Case 3 | 4 | 19 | 0 |
| Case 4 | 0 | 754 | 0 |
| Case 5 | ND<sup>b</sup> | 78 | 0 |
| Case 7 | 918 | 106,188 | 213 |

ND indicates not done.
<sup>a</sup>DNA contamination was assessed by testing the RNA samples without performing a reverse transcription step to identify background complementary DNA.
<sup>b</sup>There was insufficient RNA isolated to allow for RNA-seq.

Because the HHV-6B ORF U38 is not a spliced gene product, there is a possibility of amplification of contaminating HHV-6B genomic DNA. Although we performed a DNase treatment to mitigate contaminating DNA, we performed an RT-qPCR reaction excluding reverse transcriptase to assess for genomic HHV-6B DNA contamination. There was evidence of minimal DNA contamination in only one sample (case 7; **Table 5**).

### Host cellular gene transcriptome and cytokines in whole blood samples from post-HCT cases and controls

We performed a differential gene expression analysis of host cellular gene expression profiles in whole blood comparing normalized read counts from RNA-seq data sets between post-HCT cases and controls to determine whether we could distinguish between individuals with and without HHV-6B reactivation. We did not observe statistically significant differences in cellular gene expression in any of the paired samples (data not shown). Using a commercially available Luminex based assay, we also tested plasma samples obtained pre- and post-PAXgene tube whole blood collection for 14 immuno-inflammatory cytokines from cases and controls (see **Supplement**). All the cytokines tested in plasma were at or below the limit of detection and precluded analysis of differential cytokine expression between individuals with and without HHV-6B reactivation (data not shown).

### Structure of select HHV-6B transcripts from whole blood and DLBCL tumor tissue samples during HHV-6B replication

We recently reported differential splicing patterns of several HHV-6B transcripts in cell lines following *in vitro* infection, sometimes involving unpredicted splicing sites compared to the reference genome annotation.^26^ These findings could affect the performance of diagnostic RT-qPCR assays targeting these transcripts. In the current study, we characterized select HHV-6B spliceoforms that were detected at relatively high levels (ORFs U79, U95, and U100) in post-HCT whole blood and DLBCL tumor tissue samples. Our analysis of the mapped reads revealed that all detected splice sites contained the canonical dinucleotides GT and AG for donor and acceptor sites, respectively (**Table S3**).

For the HHV-6B U79 transcript, four different mRNA spliceoforms have been previously described and sequenced.^27^ An additional U79 mRNA isoform was recently identified in T cell lines following experimental infection with HHV-6B, and its translation into protein was confirmed using mass spectrometry.^26^ In our current study, RNA-seq data from whole blood samples demonstrated an additional novel U79 splice variant corresponding to an open reading frame including exon I, exon II, intron II, and a partial sequence from exon III (**Figure S1)**. Using proteomic data from a previously published study,^26^ we discovered a HHV-6B peptide sequence corresponding to the sequence encoded by intron II of the U79 mRNA (peptide sequence VYAQVGGVLGSPKR). This U79 mRNA isoform was also detected in one of the 3 U79 mRNA isoforms from the DLBCL tumor tissue.

The structure of the HHV-6B U95 transcript has been previously reported.^27^ Our RNA-seq data from whole blood samples confirmed that the transcription start site is located upstream of the initiating ATG sequence and that the transcript consists of two exons on both sides of ORF U94.

For the HHV-6B U100 transcript, a previous study showed the viral transcripts encode for two glycoproteins designated gQ1 and gQ2, both of which are expressed on the viral envelope in the form of a tetrameric complex with glycoproteins gH and gL.^27^ Our RNA-seq data from whole blood samples revealed reads mapping to either gQ1 or gQ2. In contrast, data from the DLBCL tumor tissue reads mapped to both gQ1 and gQ2 simultaneously. These observations suggest that each glycoprotein expression could be independently regulated, which is further supported by the existence of a polyA sequence at the end of the gQ1 open reading frame. Additionally, our analysis of reads mapping to the gQ2 transcript showed two possible splice sites, and the mature transcript corresponding to the alternative gQ2 splice is predicted to code for a shorter protein (68 amino acids versus 182 amino acids, **Figure S2**). Further experiments are required to establish the biological relevance of this gQ2 protein isoform.

## DISCUSSION

We used RNA-seq to characterize the *in vivo* HHV-6B transcriptome in whole blood samples collected from allogeneic HCT recipients and compared this to the HHV-6B transcriptome in DLBCL tumor tissue and to HHV-6B infected SupT1 CD4+ cells in culture. We observed concordance between the HHV-6B transcriptome observed in the *in vivo* and *in vitro* samples, although there was substantial variability in the breadth and quantity of gene expression across sample types. The HHV-6B viral polymerase gene U38 was the only transcript detected in all RNA-seq data sets and was one of the top ten most highly expressed genes. Using a novel RT-qPCR assay, we detected the HHV-6B U38 gene in all tested whole blood samples from patients with concurrent HHV-6B viremia, suggesting that this is a sensitive assay for HHV-6B replication. The splicing pattern for the HHV-6B U79 and U100 genes in whole blood samples demonstrated discrepancies with the predicted splice sites as annotated in the HHV-6B reference genome, and caution is required in designing PCR assays for these targets.

Quantitative PCR to detect HHV-6B genomic DNA is the standard clinical approach for diagnostic testing. However, this has important limitations in specificity given that most humans are infected with HHV-6B early in life, HHV-6B remains latent in a diverse array of cell types, and about 1% of the population has iciHHV-6 (species A or B) with a high burden of cell-associated HHV-6 DNA.^10^ Thus, it may be difficult to distinguish between latent and actively replicating virus when HHV-6B DNA is detected by qPCR in cellular samples. For a related herpesvirus, cytomegalovirus (CMV), the use of RT-qPCR to identify CMV transcripts in bronchoalveolar lavage fluid of immunocompromised patients demonstrated significantly greater specificity for CMV pneumonia than standard qPCR, which has poor specificity due to the detection of latent virus and viral shedding.^28^ Development of RT-qPCR assays to detect HHV- 6B mRNA may provide a more accurate method for identifying active infection, and studies using RT-qPCR to detect spliced transcripts have demonstrated relatively high sensitivity and specificity for viral replication *in vivo*.^11,16,17^ However, HHV-6B transcript targets that have been studied to date were selected based on their temporal expression pattern or because they were a spliced gene product.

This is the first study to our knowledge using RNA-seq to identify the HHV6-B gene expression profile in whole blood collected during periods of HHV-6B reactivation. Previous reports have characterized the expression pattern of several HHV-6B transcripts following experimental *in vitro* infection of different T cell lines with HHV-6B,^26^ and one study described the HHV-6B transcriptome in tumor tissue samples from two patients with DLBCL,^19^ although the pathogenic role of HHV-6B in this context is unclear. RNA-seq for transcriptome analysis provides an unbiased approach to identify consistently and highly expressed genes that may increase the sensitivity of targeted RT-qPCR assays for HHV-6B replication in biologic samples. Given that total mRNA isolation from PBMCs in blood samples may be limited by low yields, degradation, coagulation, and the fact that cell purification procedures can skew transcriptome analysis by inducing gene transcription,^29^ we chose to perform direct mRNA isolation from whole blood samples using PAXgene tubes to allow for immediate and long-term stabilization of intracellular RNA.^29^

We demonstrated a broad range of viral gene expression across the HHV-6B genome in whole blood from subjects with concurrent plasma viremia, indicative of lytic replication. We did not detect any HHV-6B transcripts in subjects without concurrent plasma viremia. The previously reported study of two DLBCL tumor samples with HHV-6B infection demonstrated higher RNA-seq read counts mapping to the HHV-6B genome with broader gene expression profiles including almost all known HHV-6B genes.^19^ Hierarchical cluster analysis of the HHV-6B transcriptome from whole blood versus DLBCL tumor samples with HHV-6B infection showed similarities but also differences in gene expression profiles that may be explained in part by differences in infected cell types. Nonetheless, we identified five gene transcripts that were represented among the top ten most highly expressed HHV-6B transcripts in all *in vivo* and *in vitro* samples (**Table 4)**.

HHV-6B U38 was the only gene transcript detected in all the RNA-seq data sets and was the only viral transcript detectable in two of the whole blood samples. Interestingly, HHV-6B U38 appears to be an immunodominant epitope recognized by human CD4+ T cells.^30^ We developed a targeted RT-qPCR assay for U38, which was positive in all tested whole blood samples from patients with HHV-6 viremia, demonstrating the potential utility of this assay for identifying HHV-6B replication in clinical samples. Although U38 is not a spliced gene product leading to the possibility of contamination with genomic DNA, we demonstrated minimal contamination after treatment with DNase. Whereas prior studies have tested the sensitivity and specificity of assays targeting HHV-6B transcripts such as U12, U16/17, U60/66, U79/U80, U89/90, and U100,^11–17^ U79 was the only one of these transcripts shared across sample types at a relatively high abundance. Our data also demonstrated potential unique spliceoforms for U79 and U100, which may further compromise the sensitivity of PCR assays for these targets.

We also used the RNA-seq data from whole blood to study whether unique host gene expression or cytokine profiles could be demonstrated among patients with and without HHV-6 reactivation post-HCT. Provocative data using microarrays demonstrated a unique host gene expression profile in children with primary HHV-6B infection,^31^ and other studies have shown associations between specific cytokine levels and HHV-6 infection after HCT.^32^ However, we did not find any significant differences in whole blood host gene expression and cytokine profiles between subjects with and without HHV-6 reactivation in this study. However, our sample size was small, no subjects in our cohort had severe HHV-6 infection with end-organ disease, and biologic changes early after HCT (e.g. donor cell engraftment, graft-versus-host disease) likely had larger effects on host gene expression.

Strengths of this study included the use of RNA-seq analyses of unique biologic samples to characterize *in vivo* HHV-6B gene expression and comparison of these findings with *in vitro* HHV-6B gene expression. Furthermore, we used these data to generate an evidence-based RT-qPCR assay to identify HHV-6B replication in biologic samples. The low overall number of HHV-6B gene transcript reads prevented us from accurately determining the relative abundance of each transcript. Nonetheless, our characterization of the HHV-6B transcriptome in HCT patients with HHV-6B reactivation provides a comprehensive list of viral proteins of biological interest. Future studies using sequentially collected samples from subjects before and after HHV-6B reactivation, as well as in subjects with iciHHV-6, will be important to refine these findings.

In conclusion, we demonstrated that RNA-seq can be used to determine HHV-6B gene expression after allogeneic HCT, and these analyses identified the HHV-6B U38 mRNA transcript as a potential target for focused clinical diagnostic assays such as RT-qPCR. A novel RT-qPCR assay targeting HHV-6B U38 was established and detected U38 mRNA in all tested samples from subjects with HHV-6B viremia, underscoring the potential for this biomarker to identify active viral replication in clinical samples. Larger studies that correlate HHV-6B mRNA transcript detection with HHV-6B genomic DNA detection and HHV-6B disease will be critical to establish actionable DNA and mRNA transcript thresholds for treatment.

## Authorship Contributions

JAH, MB, and SB were responsible for the design of the study. JAH, MI, RB, VP, M-LH, RHS, KRJ, and SB analyzed the data. Samples and data were collected by JAH and SB. JAH, DMZ, RB, RHS, KRJ, MB, and SB interpreted the data. JAH and SB wrote the first draft. All authors contributed to the writing and revision of the manuscript and approved the final version.

## Acknowledgements

We would like to acknowledge the Fred Hutchinson Cancer Research Center Research Cell Bank for providing B-LCL samples and the HHV-6 Foundation for providing HHV-6B infected SupT1 CD4+ T cells. We also thank Terry Stevens-Ayers, Elsa Garnace, and Zach Stednick for help obtaining samples and data.

## References

1. Hill JA, Zerr DM. Roseoloviruses in transplant recipients: clinical consequences and prospects for treatment and prevention trials. Curr Opin Virol. 2014;9:53–60. doi:10.1016/j.coviro.2014.09.006.

2. Scheurer ME, Pritchett JC, Amirian ES, Zemke NR, Lusso P, Ljungman P. HHV-6 encephalitis in umbilical cord blood transplantation: a systematic review and metaanalysis. Bone Marrow Transplant. 2013;48(4):574–580. doi:10.1038/bmt.2012.180.

3. Achour A, Boutolleau D, Slim A, Agut H, Gautheret-Dejean A. Human herpesvirus-6 (HHV-6) DNA in plasma reflects the presence of infected blood cells rather than circulating viral particles. J Clin Virol. 2007;38(4):280–285. doi:10.1016/j.jcv.2006.12.019.

4. Hill JA, Boeckh MJ, Sedlak RH, Jerome KR, Zerr DM. Human herpesvirus 6 can be detected in cerebrospinal fluid without associated symptoms after allogeneic hematopoietic cell transplantation. J Clin Virol. July 2014. doi:10.1016/j.jcv.2014.07.001.

5. Hill J a, Koo S, Guzman Suarez BB, et al. Cord-blood hematopoietic stem cell transplant confers an increased risk for human herpesvirus-6-associated acute limbic encephalitis: a cohort analysis. Biol Blood Marrow Transplant. 2012;18(11):1638–1648. doi:10.1016/j.bbmt.2012.04.016.

6. Fotheringham J, Akhyani N, Vortmeyer A, et al. Detection of active human herpesvirus-6 infection in the brain: correlation with polymerase chain reaction detection in cerebrospinal fluid. J Infect Dis. 2007;195(3):450–454. doi:10.1086/510757.

7. Nagate et al. Detection and quantification of human herpesvirus 6 genomes using bronchoalveolar lavage fluid in immunocompromised patients with interstitial pneumonia. Int J Mol Med. 2001;8(4):379.

8. Kaufer BB, Flamand L. Chromosomally integrated HHV-6:impact on virus, cell and organismal biology. Curr Opin Virol. 2014;9C:111–118. doi:10.1016/j.coviro.2014.09.010.

9. Hall Sedlak R, Hill JA, Nguyen T, et al. Detection of HHV-6B reactivation in hematopoietic cell transplant recipients with inherited chromosomally integrated HHV-6A by droplet digital PCR. J Clin Microbiol. February 2016. doi:10.1128/JCM.03275-15.

10. Hill JA, Hall Sedlak R, Jerome KR. Past, Present, and Future Perspectives on the Diagnosis of Roseolovirus Infections. Curr Opin Virol. 2014:in press.

11. Norton RA, Caserta MT, Hall CB, Schnabel K, Hocknell P, Dewhurst S. Detection of human herpesvirus 6 by reverse transcription-PCR. J Clin Microbiol. 1999;37(11):3672–3675.

12. Van den Bosch G, Locatelli G, Geerts L, et al. Development of reverse transcriptase PCR assays for detection of active human herpesvirus 6 infection. J Clin Microbiol. 2001;39(6):2308–2310. doi:10.1128/JCM.39.6.2308-2310.2001.

13. Kondo K, Kondo T, Shimada K, Amo K, Miyagawa H, Yamanishi K. Strong interaction between human herpesvirus 6 and peripheral blood monocytes/macrophages during acute infection. J Med Virol. 2002;67(3):364–369. doi:10.1002/jmv.10082.

14. Pradeau K, Bordessoule D, Szelag J-C, et al. A reverse transcription-nested PCR assay for HHV-6 mRNA early transcript detection after transplantation. J Virol Methods. 2006;134(1-2):41–47. doi:10.1016/j.jviromet.2005.11.015.

15. Strenger V, Caselli E, Lautenschlager I, et al. Detection of HHV-6-specific mRNA and antigens in PBMCs of individuals with chromosomally integrated HHV-6 (ciHHV-6). Clin Microbiol Infect. 2014;20(10):1027–1032. doi:10.1111/1469-0691.12639.

16. Ihira M, Enomoto Y, Kawamura Y, et al. Development of quantitative RT-PCR assays for detection of three classes of HHV-6B gene transcripts. J Med Virol. 2012;84(9):1388–1395. doi:10.1002/jmv.23350.

17. Bressollette-Bodin C, Nguyen TVH, Illiaquer M, et al. Quantification of two viral transcripts by real time PCR to investigate human herpesvirus type 6 active infection. J Clin Virol. 2014;59(2):94–99. doi:10.1016/j.jcv.2013.11.014.

18. Tsao EH, Kellam P, Sin CSY, Rasaiyaah J, Griffiths PD, Clark DA. Microarray-based determination of the lytic cascade of human herpesvirus 6B. J Gen Virol. 2009;90(Pt 11):2581–2591. doi:10.1099/vir.0.012815-0.

19. Strong MJ, O’Grady T, Lin Z, et al. Epstein-Barr virus and human herpesvirus 6 detection in a non-Hodgkin’s diffuse large B-cell lymphoma cohort by using RNA sequencing. J Virol. 2013;87(23):13059–13062. doi:10.1128/JVI.02380-13.

20. Hill JA, Magaret AS, Hall-Sedlak R, et al. Outcomes of hematopoietic cell transplantation using donors or recipients with inherited chromosomally integrated HHV-6. Blood. 2017;130(8). doi:10.1182/blood-2017-03-775759.

21. Zerr DM, Gupta D, Huang M-L, Carter R, Corey L. Effect of antivirals on human herpesvirus 6 replication in hematopoietic stem cell transplant recipients. Clin Infect Dis. 2002;34(3):309–317. doi:10.1086/338044.

22. Trapnell C, Pachter L, Salzberg SL. TopHat: discovering splice junctions with RNA-Seq. Bioinformatics. 2009;25(9):1105–1111. doi:10.1093/bioinformatics/btp120.

23. Langmead B, Salzberg SL. Fast gapped-read alignment with Bowtie 2. Nat Methods. 2012;9(4):357–359. doi:10.1038/nmeth.1923.

24. Anders S, Pyl PT, Huber W. HTSeq A Python Framework to Work with High-Throughput Sequencing Data. Cold Spring Harbor Labs Journals; 2014. doi:10.1101/002824.

25. Love MI, Huber W, Anders S. Moderated estimation of fold change and dispersion for RNA-seq data with DESeq2. Genome Biol. 2014;15(12):550. doi:10.1186/s13059-014- 0550-8.

26. Greninger AL, Knudsen GM, Roychoudhury P, et al. Comparative genomic, transcriptomic, and proteomic reannotation of human herpesvirus 6. BMC Genomics. 2018;19(1):204. doi:10.1186/s12864-018-4604-2.

27. Taniguchi T, Shimamoto T, Isegawa Y, Kondo K, Yamanishi K. Structure of Transcripts and Proteins Encoded by U79-80 of Human Herpesvirus 6 and Its Subcellular Localization in Infected Cells. Virology. 2000;271(2):307–320. doi:10.1006/viro.2000.0326.

28. Boivin G, Olson CA, Quirk MR, Kringstad B, Hertz MI, Jordan MC. Quantitation of cytomegalovirus DNA and characterization of viral gene expression in bronchoalveolar cells of infected patients with and without pneumonitis. J Infect Dis. 1996;173(6):1304–1312.

29. Rainen L, Oelmueller U, Jurgensen S, et al. Stabilization of mRNA expression in whole blood samples. Clin Chem. 2002;48(11):1883–1890.

30. Nastke M-D, Becerra A, Yin L, et al. Human CD4+ T cell response to human herpesvirus 6. J Virol. 2012;86(9):4776–4792. doi:10.1128/JVI.06573-11.

31. Hu X, Yu J, Crosby SD, Storch GA. Gene expression profiles in febrile children with defined viral and bacterial infection. Proc Natl Acad Sci U S A. 2013;110(31):12792– 12797. doi:10.1073/pnas.1302968110.

32. Ogata M, Satou T, Kawano R, et al. Correlations of HHV-6 viral load and plasma IL-6 concentration with HHV-6 encephalitis in allogeneic stem cell transplant recipients. Bone Marrow Transplant. 2010;45(1):129–136. doi:10.1038/bmt.2009.116.

33. Øster B, Höllsberg P. Viral gene expression patterns in human herpesvirus 6B-infected T cells. J Virol. 2002;76(15):7578–7586. doi:10.1128/JVI.76.15.7578-7586.2002.

